# Multiplexed encoding of frequency-modulated sweep features in the inferior colliculus

**DOI:** 10.1101/2025.02.10.637492

**Authors:** Audrey C. Drotos, Sarah Z. Wajdi, Michael Malina, Marina A. Silveira, Ross S. Williamson, Michael T. Roberts

## Abstract

Within the central auditory pathway, the inferior colliculus (IC) is a critical integration center for ascending sound information. While IC neurons have well-characterized receptive fields for individual sound features such as sound frequency, intensity, and location, growing evidence suggests that some neurons also use multiplexing to encode sound feature combinations. Here, we performed in vivo juxtacellular recordings in awake, head-fixed mice to examine how individual IC neurons and neuronal populations encode the speed, direction, and frequency range of frequency-modulated sweeps. To understand the strategies used by neurons to represent different sound features, we trained a support vector machine to decode sound features from different parameters of the spike train, including the firing rate, spike times relative to stimulus onset, distribution of inter-spike intervals, and first spike latency. We found that many IC neurons multiplex features of frequency-modulated (FM) sweeps using distinct temporal coding strategies rather than simple changes in mean firing rate, and that these feature representations are interdependent, yielding a combinatorial encoding of sound features within individual neurons. Accordingly, using static receptive fields for sweep frequency or direction alone yielded poor predictions of neuron responses to vocalizations that contain simple frequency changes. Lastly, we showed that encoding strategies varied across individual neurons, resulting in a highly informative population code for FM sweep parameters. Together, our results suggest that multiplexing is a common mechanism used by IC neurons to represent complex sound features.

## Introduction

One of the primary functions of the brain is to build representations of complex sensory objects based on the relatively limited set of features encoded by the sensory periphery. Many previous studies have shown that neurons in the inferior colliculus (IC), the midbrain hub of the central auditory pathway, are tuned to specific sound features. For example, IC neuron firing rate can be modulated by the frequency and intensity of pure tone stimuli (Suga, 1964; Ramachandran et al., 1999; Palmer et al., 2013), and IC neurons can exhibit firing-rate based tuning for sound duration (Casseday et al., 1994; Ehrlich et al., 1997; Fuzessery and Hall, 1999; Pérez-González et al., 2006) and sound location (Aitkin et al., 1984; Kuwada et al., 1987; Davis et al., 2003; Chase, 2005; Slee and Young, 2011). Prior studies also suggest that some IC neurons use changes in firing rate to encode the direction of rapid movements across sound frequency known as frequency-modulated (FM) sweeps (Suga, 1965; Poon et al., 1991; Fuzessery, 1994; Fuzessery et al., 2006; Andoni et al., 2007; Xie et al., 2007; Gittelman et al., 2009; Kuo and Wu, 2012; Geis and Borst, 2013).

However, individual neurons have many available strategies by which they can encode sound features. In addition to firing rate, neurons can use the temporal patterning of spikes, including the timing of spikes over the duration of the stimulus, the inter-spike interval distributions, and/or first spike latency to transmit information about stimuli to postsynaptic partners. Neurons can also use combinations of these strategies to encode more than one stimulus feature in a single spike train, a coding approach called multiplexing. Evidence for multiplexing in auditory neurons has been found in auditory cortex (Furukawa and Middlebrooks, 2002; Lakatos et al., 2007; Walker et al., 2011; Insanally et al., 2019), and growing evidence suggests it also exists in IC (Zheng and Escabí, 2008; Day and Delgutte, 2016; Caruso et al., 2018). For example, the spiking patterns of single IC neurons carry information about three sound localization cues (Chase and Young, 2008), and IC neurons can also co-encode information about sound and task-relevant variables in a behavioral task (Quass et al., 2024). In addition, IC neurons can simultaneously encode information across sensory modalities, including visual and auditory stimuli (Schmehl et al., 2025).

Multiplexing sound features within single IC neurons requires the use of multiple coding strategies. While most previous studies have focused on rate coding as the primary way that IC neurons convey information about sound features, temporal codes also play a strong role in the IC in the form of first-spike latencies (Chase and Young, 2008; Zheng and Escabí, 2008; Chot and Zhang, 2025) and phase-locking to stimulus features (Joris et al., 2004; Nelson and Carney, 2007; Cai and Dent, 2020). Emerging evidence suggests that temporal codes are also important in higher-order auditory structures such as auditory cortex (Bagur et al., 2025). This raises the possibility that IC neurons could use combinations of changes in spike timing and spike rate to multiplex features of complex sounds, but whether and how IC neurons use such coding strategies is not well understood.

To examine this, we performed in vivo, juxtacellular recordings from IC neurons in awake mice during presentations of frequency-modulated (FM) sweeps and used neural decoding to examine how sweep features are encoded within spike trains. We found that FM sweep features including sweep direction, speed, and frequency range could be decoded from the mean firing rates, time-binned spike counts, ISI distributions, and/or first spike latencies in individual neurons, revealing that IC neurons use many strategies to encode stimulus-specific information. In addition, neuron responses to simple sound features such as sweep direction did not predict responses to complex sounds such as vocalizations, indicating that complex sound responses are not a simple extension of neuron tuning properties. Overall, these heterogeneous sound responses in individual IC neurons yielded a population code that robustly represented complex sound features. Thus, our results show that many IC neurons use multiplexed coding strategies and that multiplexing at the individual neuron level can yield effective population codes for complex auditory stimuli.

## Results

### IC neurons in awake mice use spike rate and timing to encode the direction of four-octave FM sweeps

Many previous studies quantified direction selectivity for FM sweeps by determining the sweep direction (up or down) that elicits a larger number of spikes (Fuzessery et al., 2006; Gittelman et al., 2009; Kuo and Wu, 2012; Geis and Borst, 2013; Morrison et al., 2018), ignoring the roles that spike timing might play in encoding information about sweep identity. To overcome this problem, we performed in vivo juxtacellular recordings from IC neurons in awake, head-fixed mice (**Figure 1A-B**) and examined direction encoding using both an asymmetry index comparing the number of spikes elicited in each sweep direction and a machine learning model that was trained on the time-binned spike counts for each trial. Recordings covered the tonotopic map of the IC with neuron best frequencies (the frequency eliciting the greatest number of spikes at 70 dB SPL) ranging from 4-64 kHz (**Figure 1C**). To investigate direction selectivity, we played 70 dB SPL logarithmic FM sweeps with speeds of 10 – 200 octaves/second. Neurons exhibited a diversity of responses, including direction selectivity (**Figure 1D-F**, first and third cell), asymmetries in spike timing between up and down sweeps (**Figure 1D-F**, second cell), and inhibition (**Figure 1D-F**, fourth cell). We first evaluated direction selectivity for neurons in our dataset using the direction selectivity index (DSI), which is a normalized comparison of the firing rates of the neuron during the up sweep compared to the down sweep (see equation in Methods). Neurons with a DSI closer to +1 have higher firing rates in the upward sweep direction, while neurons with a DSI closer to −1 have higher firing rates in the downward direction (Kuo and Wu, 2012). We found that the DSIs of the cells we recorded from formed a continuum (**Figure 1I**), with most cells being non-selective according to the criteria of Kuo and Wu, 2012 (37/46 cells, defined as DSIs < +0.33 and > −0.33). Of the 9 cells that were directionally selective according to the DSI metric, 4 were selective for downward sweeps and 5 were selective for upward sweeps.

**Figure 1.**
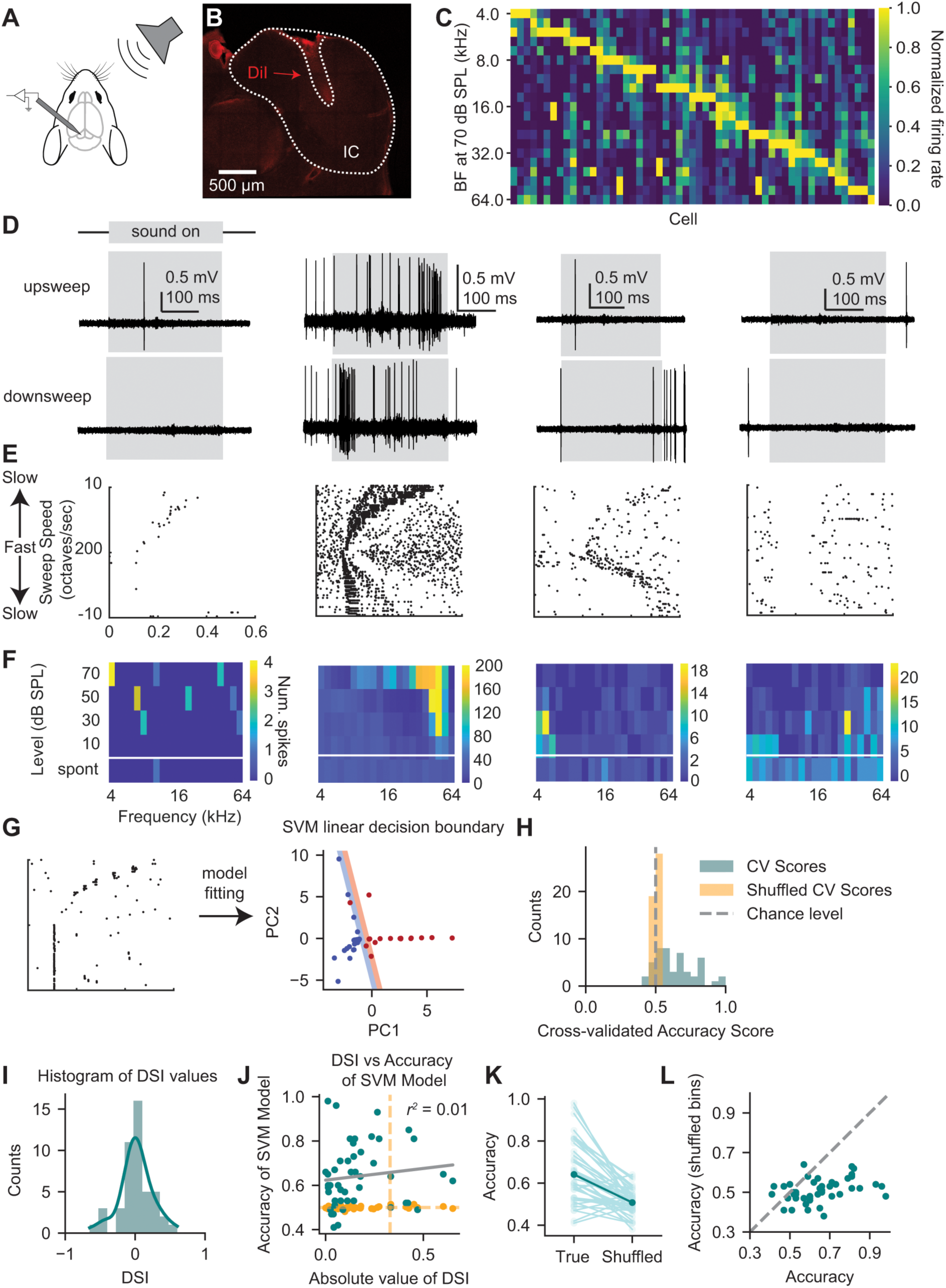
IC neurons use spike timing to encode direction information for FM sweeps. *A,* Experimental design. Awake mice were headfixed in a sound booth with sound presented to the right side of the mouse while juxtacellular recordings were performed in the left IC. ***B,*** Example of a DiI labeled recording site in the left IC, outlined in white. ***C,*** The tonal receptive field at 70 dB SPL for each neuron included in the study, sorted by best frequency. ***D,*** Four example neuron responses to the slowest (10 octaves/s) up (top row) and down (bottom row) frequency-modulated (FM) sweeps. Grey boxes indicate when the sound was presented. ***E,*** Raster plots showing responses to four-octave FM sweeps for neurons shown in *D*. The top half of the plot shows responses to upward sweeps, and the bottom half of the plot shows responses to downward sweeps. Sweep speed varies along the y-axis with the fastest sweep speed shown at the middle of the y-axis (200 octaves/s) and sweep speeds becoming progressively slower toward the top and bottom of the y-axis. ***F,*** Frequency response areas (FRAs) for neurons shown in *D.* FRAs were constructed using 200 ms tone pips presented in pseudorandom order to cover a 4-64 kHz frequency range at 5 steps/octave intervals and the listed intensities. The bottom row of the heatmap indicates the spontaneous firing rate of the neuron. ***G,*** Example of a hyperplane from a model trained to decode direction from neural spike time data. For visualization purposes, data were binned into 16 bins and then PCA-transformed and projected onto the first two principal components. Colored regions on each side of the hyperplane indicate the margins. ***H,*** Direction decoding accuracy scores for 4-64 kHz FM sweeps. Blue bars indicate accuracy scores for model trained on true data; orange bars indicate accuracy for model trained on data with shuffled class labels. ***I,*** Histogram of direction selectivity indices (DSIs) calculated from the same group of cells shown in *H.* The histogram is overlaid with a kernel density estimate that was generated using the ‘kde’ argument for seaborn’s ‘histplot’ function. ***J,*** Scatterplot comparing the accuracy of the model for decoding direction and the absolute value of the DSI in individual neurons. Blue dots are true accuracy, orange dots are accuracy of the model with shuffled class labels. Grey line shows the linear regression between true accuracy labels and DSI, indicating no correlation between the two values. Vertical orange line is the DSI cutoff from Kuo and Wu (2012); neurons to the right of this line are considered direction selective according to the DSI metric. Horizontal orange line is model chance. ***K,*** Comparison of model accuracy for direction decoding with time bins in correct order vs. shuffled time bins. Class labels are maintained in each case. Individual neurons are connected by lines, and the mean is shown in dark blue. ***L,*** Scatterplot showing the accuracy for true bin order vs. shuffled bin order (same as *K*). The dashed grey line indicates the line of identity (x = y).

However, the DSI measure only considers the total number of spikes occurring during the up or down sweep, disregarding information about spike timing that could be used by downstream neurons to decode direction. To overcome this problem, we used a linear support vector machine (SVM), a supervised machine learning approach that classifies data points by fitting linear hyperplanes that separate classes (sweep directions) in a high-dimensional space. We trained an SVM on the time-binned spike counts from up and down sweep trials to investigate whether the direction of the sweep could be decoded from the spike trains of individual IC neurons (**Figure 1G**, example shown uses 16 time bins principal components analysis (PCA) transformed and projected onto the first two principal components for illustrative purposes). Using the model, we found that the direction of the FM sweep could be determined significantly above chance in 25/46 neurons (**Figure 1H**). Interestingly, we found that the group of neurons where direction could be significantly decoded using the model did not completely overlap with the group of neurons that were deemed directionally selective using the DSI. Specifically, we found that sweep direction could be decoded from 20 cells that had non-selective DSIs and we found no significant correlation (r^2^ = 0.01) between the absolute value of neuron DSI and model accuracy (**Figure 1J**).

We hypothesized that the mismatch between the DSI and model results occurred in part because the timing of spikes, which is present within the time-binned spike counts and accessible to the SVM but is not used by the DSI, contains information about sweep direction. To test this hypothesis, we shuffled the order of the time bins within the time-binned spike counts while maintaining class labels and compared the accuracy of the shuffled model to the true accuracy (bin order intact) for each neuron. We found a significant decrease in the model accuracy across all the cells when the time bins were shuffled (paired t-test, *t* = 7.0, *p* = 1.0e-8). There were only 2 cells from which the direction could be decoded with shuffled bins compared to 25 when the time bin order was intact, suggesting that the timing of spikes, rather than just the total number of spikes, is critical for encoding direction in individual neurons (**Figure 1K**). In particular, we found that cells that more accurately encoded direction had the largest shifts in accuracy after the time bins were shuffled (**Figure 1K,L**), suggesting that cells that encode direction well are more likely to use temporal codes. Overall, these results suggest that neurons use a combination of strategies including both the timing of individual spikes and the overall firing rate to encode sweep direction for FM sweeps.

### Direction encoding for four-octave FM sweeps varies across sweep intensities but not speed

We next investigated how neural tuning for sweep direction is affected by changes in sweep speed and sweep intensity. We hypothesized that sweep decoding would be highest at slower sweep speeds, given this would exaggerate the asynchronies in activation of excitatory and inhibitory areas of a neuron’s tonal receptive field that have been suggested to underlie sweep selectivity (Xie et al., 2007; Kuo and Wu, 2012). To examine this, we played upward and downward four-octave FM sweeps with speeds varying from 10 – 200 octaves/second at four different sound intensities (10, 30, 50, and 70 dB SPL, **Figure 2A**) and used an SVM to decode the sweep direction at each intensity level and each speed. In line with our hypothesis, we found that there was a significant difference in direction decoding across speeds (**Figure 2B**, repeated-measures ANOVA, *F*(9, 414) = 2.5, *p* = 0.0077). However, post-hoc tests revealed that none of the pairwise accuracies were significantly different from each other (Tukey’s HSD, **Figure 2C**). In addition, while the number of cells from which direction could be decoded increased as speed decreased, this trend was also not significant (Chi-square test for trend, χ*^2^* = 3,69, *p* = 0.93). These results suggest that some IC neurons can robustly encode the direction of four-octave FM sweeps across the tested range of sweep speeds.

**Figure 2.**
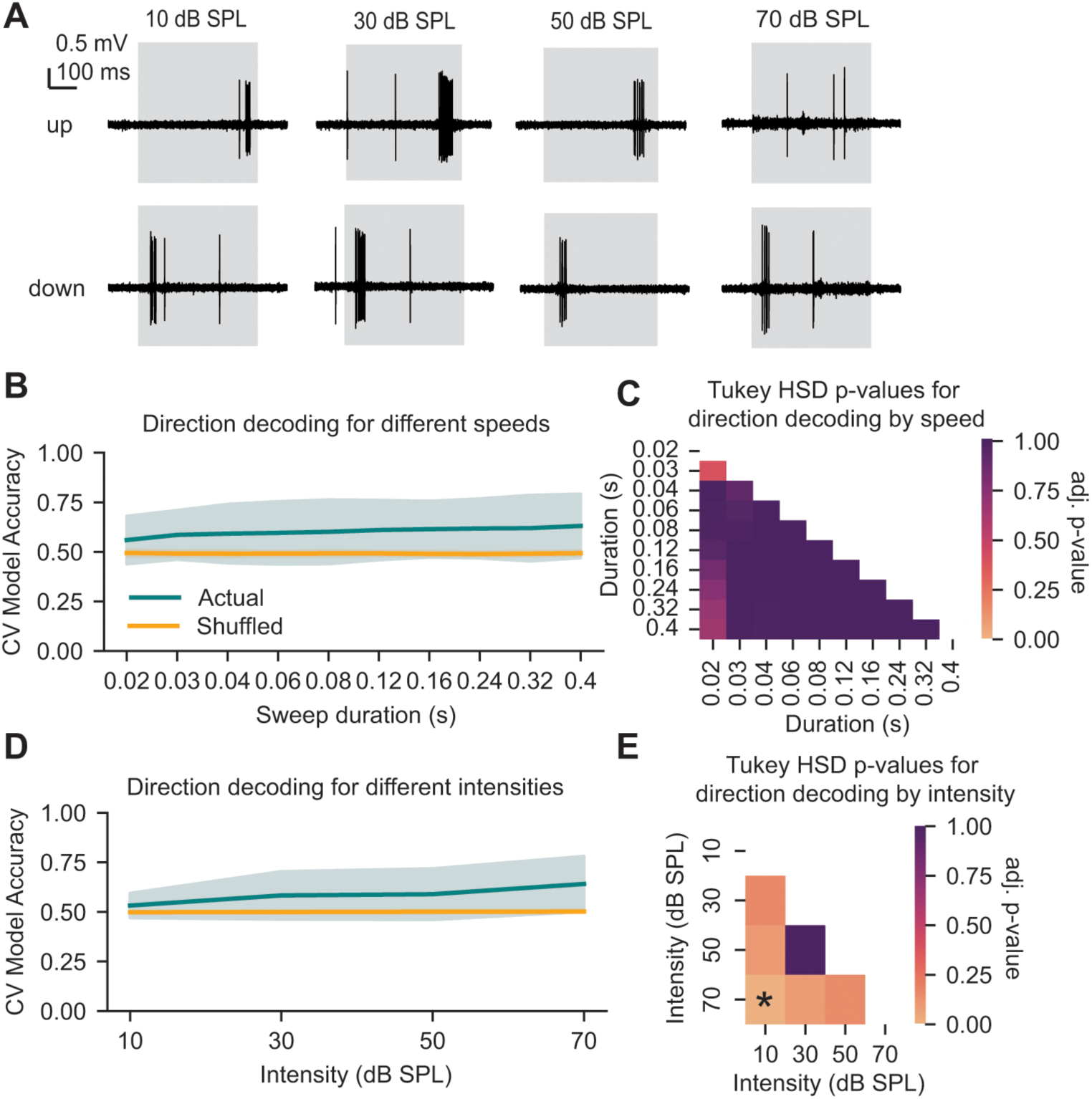
Four-octave FM sweep direction decoding is consistent across varying speeds but not intensities. *A,* Sample traces from a recording showing differing responses to four-octave up and down sweeps (10 octaves/second) at each intensity level. Sound presentation is indicated by the grey box. ***B,*** Accuracy of direction decoding across varying sweep speeds. The blue line indicates direction decoding accuracy across speeds for true labels, and the orange line indicates accuracy for shuffled labels (chance). The standard deviation is shown with shading. ***C,*** Heatmap showing Tukey HSD p-values for each pairwise comparison of true data accuracy scores for each speed in *B.* Lighter colors indicate more significant results. No comparisons were statistically significant. ***D,*** Accuracy of direction decoding across varying sweep intensities. The blue line indicates direction decoding accuracy across intensities for true labels, and the orange line indicates accuracy for shuffled labels (chance). The standard deviation is shown with shading. ***E,*** Heatmap showing Tukey’s HSD p-values for each pairwise comparison of true data accuracy scores for each speed in *D.* Lighter colors indicate more significant results. Decoding accuracy at 70 dB SPL was significantly higher than accuracy at 10 dB SPL (Tukey’s HSD, *p* = 0.0001); this is indicated with an asterisk.

Next, we tested whether direction decoding accuracy was also robust across FM sweeps with varying intensities. We hypothesized that decoding accuracy would be highest at the loudest intensity tested (70 dB SPL), as this would again exaggerate spiking differences between inhibitory and excitatory areas in a neuron’s receptive field and lead to greater asymmetries in spike timing between up and down sweeps. In addition, IC neuron tonal receptive field shapes exhibit intensity-dependent changes in frequency tuning (Palmer et al., 2013), suggesting that direction selectivity may also be intensity-dependent. In line with this, we found that direction decoding accuracy in IC neurons differed across varying sweep intensities within individual neurons (**Figure 2D**, repeated-measures ANOVA, *F*(3, 138) = 12.38, *p* < 0.0001), with decoding accuracy at 70 dB SPL significantly higher than decoding accuracy at 10 dB SPL (Tukey’s HSD, *p* = 0.0001, **Figure 2E**). In addition, we observed that the number of cells from which direction could be decoded increased as intensity increased (10 dB SPL: 5/46 cells, 30 dB SPL: 14/46 cells, 50 dB SPL: 16/46 cells, 70 dB SPL: 27/46 cells), and this trend was significant (Chi-square test for trend, χ^2^ = 11.97, *p* = 0.007).

Overall, our data show that direction decoding accuracy for four-octave FM sweeps is relatively stable across sweep speed but not sweep intensity, with cells exhibiting higher decoding accuracies at higher intensities.

### Individual IC neurons robustly encode frequency range and speed but not direction of FM sweeps across varying sweep frequency ranges

Our results support the idea that some IC neurons encode sweep direction for four-octave FM sweeps. However, natural sounds rarely contain FM sweeps with such large frequency changes: many ethologically relevant sounds such as vocalizations have small frequency changes on the order of ½ to 1 octave (Holy and Guo, 2005; Portfors, 2007). Whether the direction of frequency change is encoded in IC neurons for smaller sweeps has been less well studied. To examine this, we presented both one- and two-octave FM sweeps at speeds of 10 – 200 octaves/second while recording from IC neurons in awake mice. One-octave FM sweeps consisted of the frequency ranges 4-8, 8-16, 16-32, and 32-64 kHz, and two-octave FM sweeps consisted of the frequency ranges 4-16, 8-32, and 16-64 kHz (**Figure 3A,B**).

**Figure 3.**
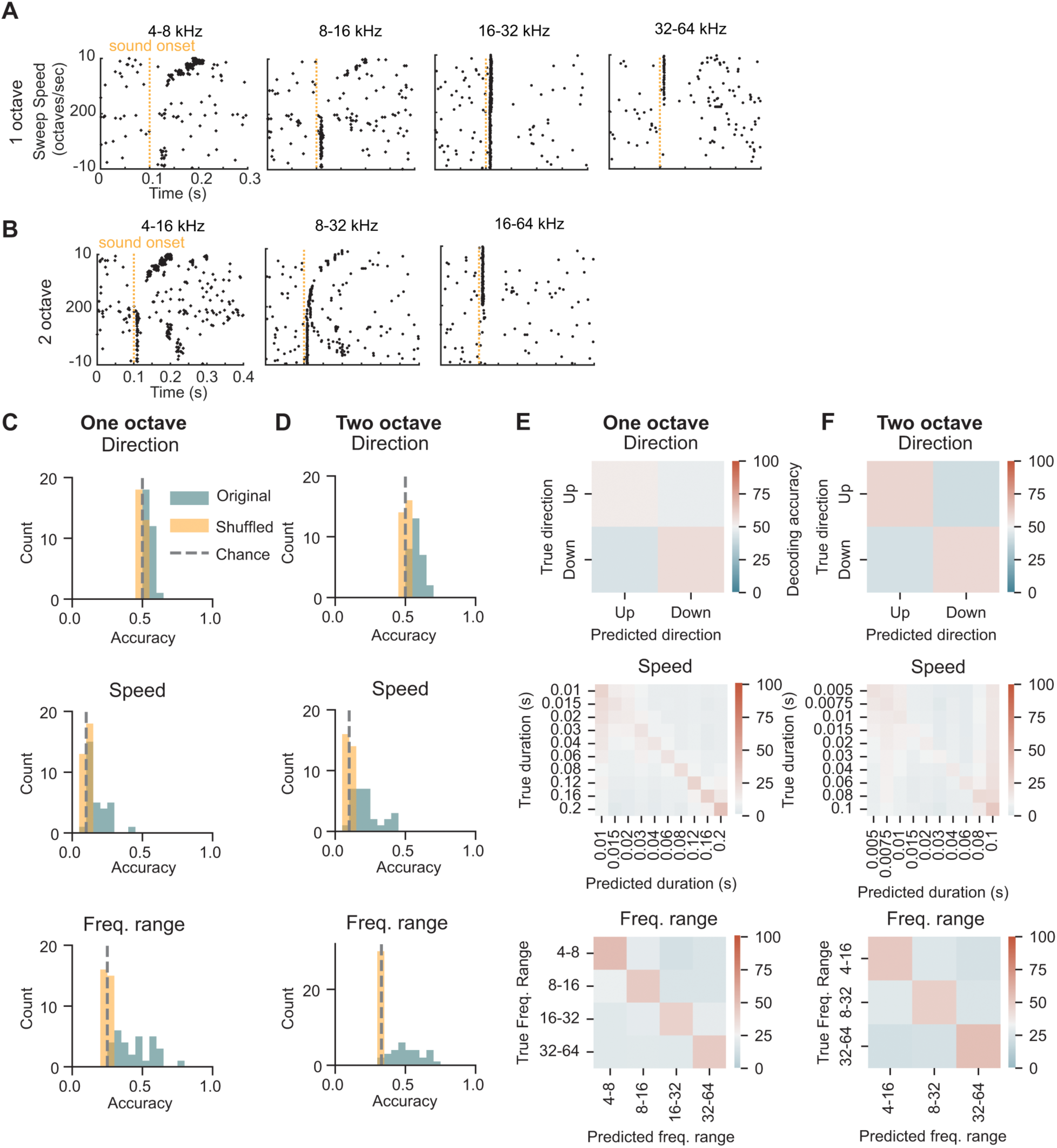
IC neurons encode frequency range, speed of one- and two-octave FM sweeps. *A,* Raster plots with responses to one-octave FM sweeps of varying sweep frequency ranges (listed on top of each plot). Up sweeps occupy the top half of the plot and down sweeps occupy the bottom. Slowest speeds are on the outsides with fastest speeds in the middle. Sound onset is shown by the orange line at 100 ms. ***B,*** Raster plots showing responses from the same neuron in *A* to the two-octave FM sweeps, played at the same speeds. ***C,*** Direction, speed, and frequency range decoding accuracy for one-octave FM sweeps. True accuracy is shown in blue, accuracy with shuffled bins is in orange. Chance for each plot is shown with a grey dashed line. ***D,*** Decoding accuracies for the same sound features shown in *C,* for two-octave FM sweeps. ***E,*** Model predictions (x-axis) versus actual (y-axis) categories for decoding of each sound feature. Predictions are aggregated across all neurons in the dataset. Chance performance is white, while above chance is red and below chance is blue. ***F-G,*** Same as A, for speed and frequency range decoding for one-octave sweeps. ***H-J,*** Same as *E*-*G* for decoding from two-octave sweeps.

To determine whether neurons were directionally selective, we used an SVM to decode the direction of the sweep from the time-binned spike counts of each neuron. Strikingly, we found that we could only decode direction from 2/31 of the cells in the one-octave sweep condition (**Figure 3C**, top), and in the two-octave condition, direction could only be decoded in 9/30 cells (**Figure 3D**, top). This finding differed from the four-octave sweep results, where direction was encoded in 25/46 cells, and suggests that selectivity for direction is not categorically encoded in individual mouse IC neurons for ethologically relevant FM sweep frequency changes.

Instead, we next asked whether other features of the sweep—in particular, the sweep frequency range and the sweep speed—could be decoded from each neuron’s response. In contrast to the direction decoding results, we found that the speed of each sweep could be decoded significantly above chance from most cells in both the one- and two-octave sweep conditions regardless of sweep direction or frequency range (25/31 and 27/30 cells, respectively, **Figure 3C,D** middle). In addition, we found that the frequency range of the sweep, regardless of sweep direction or speed, could be decoded above chance in most cells in our dataset (31/31 cells for one-octave sweeps and 24/30 cells for two-octave sweeps, **Figure 3C,D** bottom).

Prediction errors made by the model varied for all three sweep features tested. Decoding accuracy for the direction of one- and two-octave sweeps was near chance performance, with the model predicting the sweep direction incorrectly almost as often as it predicted direction correctly (**Figure 3E,F top**). Interestingly, for the one-octave sweeps the model skewed towards predicting slower sweep speeds, with lower relative accuracy for fast speeds (**Figure 3E, middle**), and the model tended to predict slower speeds than the actual stimuli for most trials. This contrasted to prediction errors for the two-octave data, where the model had highest accuracy in predicting both the fastest and the slowest sweep speeds, but tended to predict the fastest speed even for trials that were slower (**Figure 3F, middle**). Sweep frequency range decoding was similarly high across all sweep ranges tested (**Figure 3E,F bottom**). Our data suggest that individual neurons do not categorically encode the direction of one- and two-octave FM sweeps but instead provide information about other features of the stimuli, such as the frequency range and speed.

### Individual IC neurons multiplex features of FM sweeps

Previous literature suggests that individual IC neurons can jointly encode multiple sound features. For example, IC neurons can multiplex information from multiple sound localization cues with different features of a single spike train (Pena and Konishi, 2002; Chase and Young, 2008). IC neurons can also simultaneously encode visual and auditory information about a sensory stimulus (Schmehl et al., 2025), and IC neurons show selectivity for both direction and velocity for large FM sweeps (Andoni et al., 2007). We asked whether IC neurons that significantly encode one sweep feature also significantly encode other features of a sweep by correlating the decoding accuracies for direction, speed, and frequency range for each cell in our one-octave (**Figure 4A**) and two-octave (**Figure 4B**) sweep data sets. We did not find a significant positive correlation between direction and speed accuracies in the one- or two-octave data (α corrected for 6 total comparisons using the Bonferroni correction = 0.0083; linear regression, one octave: *r^2^* = 0.21, *p* = 0.019, two octave: *r^2^* = 0.20, *p* = 0.015). However, direction and frequency range decoding accuracies were strongly correlated in both the one and two octave datasets (one octave: *r^2^* = 0.65, *p* = 5.2e-8; two octave: *r^2^* = 0.69, *p* = 1.8e-8), as were frequency range and speed decoding accuracies (one octave: *r^2^* = 0.40, *p* = 0.0001; two octave: *r^2^* = 0.40, *p* = 0.0001).

**Figure 4.**
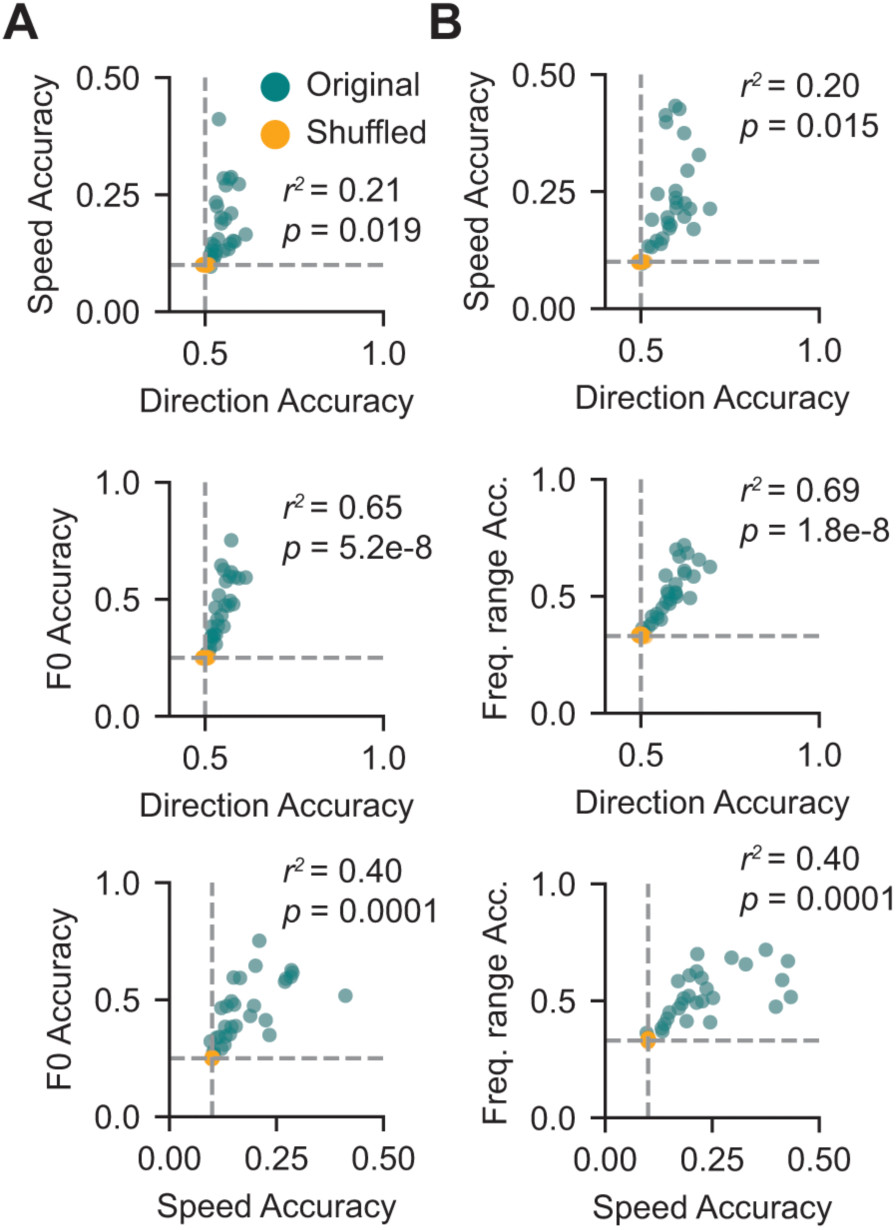
IC neurons multiplex features of one- and two-octave FM sweeps. *A,* Scatterplots showing correlations between feature encoding accuracies in individual cells for one-octave FM sweeps. Models trained on true class labels are shown in blue and models trained on shuffled class labels are shown in orange. R-squared and p-values from linear regressions between true data points for feature pairs are shown on each plot. For six total comparisons, the Bonferroni-corrected significance level (α) = 0.0083. ***B,*** Correlations of feature decoding accuracies in individual cells as in *A* for two-octave FM sweeps.

These results suggest that many IC neurons can simultaneously encode multiple FM sweep features, such as the frequency range and speed. Multiplexing features of an FM sweep would require a neuron to use different strategies to represent different features within a single spike train. There is previous evidence for multiplexing in auditory neurons: for example, in cat IC (Chase and Young, 2008) and cat auditory cortex (Furukawa and Middlebrooks, 2002) sound localization cues are encoded using multiple independent spike train features, including first spike latencies and firing rate. Some evidence for multiplexing the features of large FM sweeps also exists, with IC neurons showing selectivity for the direction and velocity of spectral motion in bat (Andoni et al., 2007). However, the specific strategies used by IC neurons to multiplex FM sweep features are not well understood.

One possibility is that in addition to overall firing rate, IC neurons use temporal dimensions of their spike trains to encode sweep features. For example, the distribution of inter-spike intervals (ISIs) can encode information in neurons independent of the temporal firing rate patterns (Bialek et al., 1991; Panzeri et al., 2001; Lundstrom and Fairhall, 2006; Insanally et al., 2019). The first spike latency (FSL) after stimulus onset can also carry important information in auditory neurons, including in cochlear nucleus (Kitzes et al., 1978) and auditory cortex (Phillips, 1988; Heil, 1997). We investigated which temporal strategies IC neurons use to encode FM sweep features by training separate decoders to predict sweep direction, frequency range, and speed from three different spike train representations: the time-binned spike counts (as shown previously), the ISI distributions, or the FSL. Interestingly, decoding accuracy scores obtained using each strategy were not significantly different from each other across the population for decoding direction (Friedman test, one octave: χ^2^(30) = 1.03, *p* = 0.60, **Figure 5A**, two octave: χ^2^(29) = 0.07, *p* = 0.97, **Figure 5G**), speed (one octave: χ^2^(30) = 0.06, *p* = 0.96, **Figure 5B**, two octave: χ^2^(29) = 2.87, *p* = 0.24, **Figure 5H**) or frequency range (one octave: χ^2^(30) = 0.58, *p* = 0.75, **Figure 5C**, two octave: χ^2^(29) = 3.20, *p* = 0.20, **Figure 5I**).

**Figure 5.**
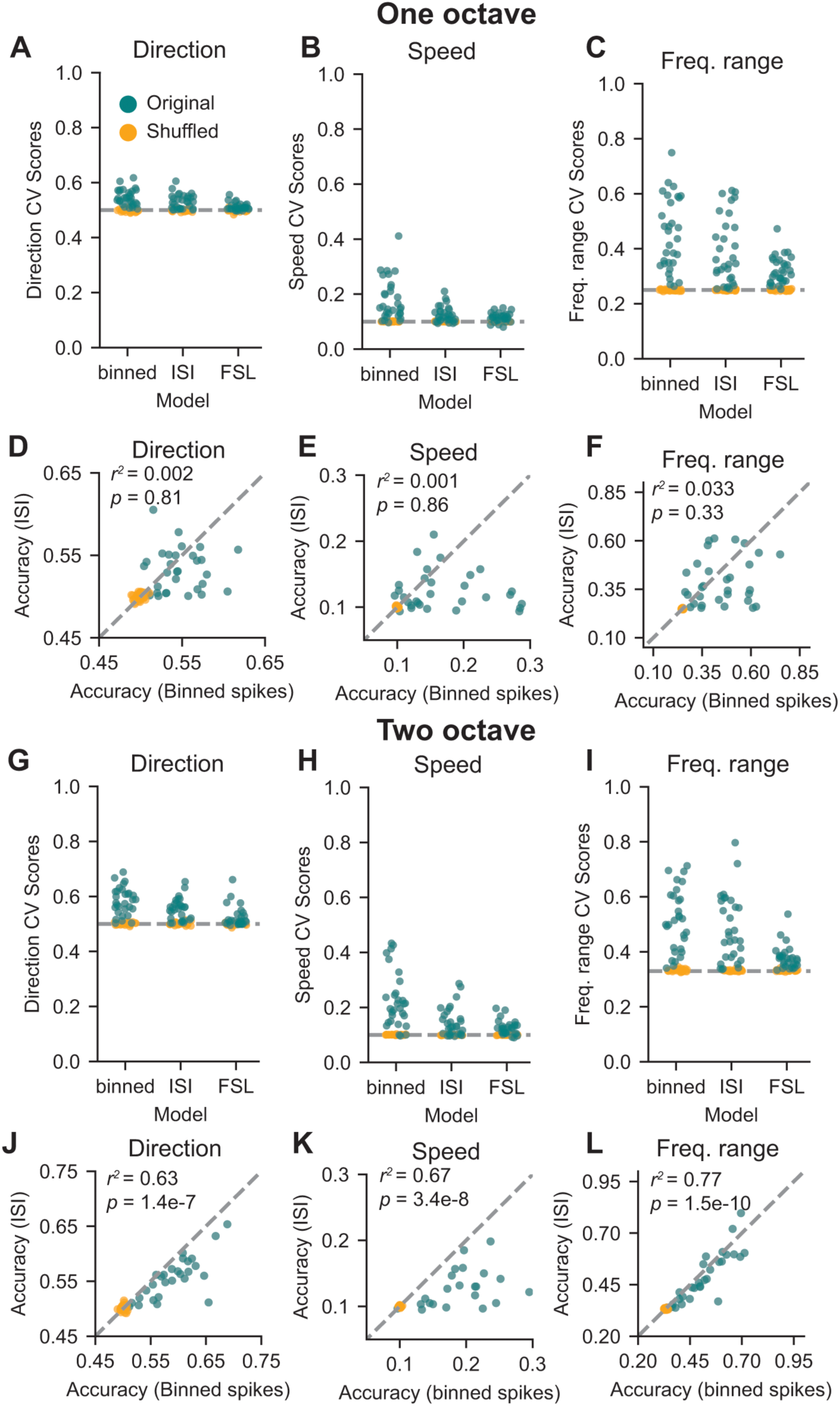
IC neurons use multiple strategies to encode FM sweep features in a single spike train. *A,* Accuracies for decoding direction in one-octave sweeps from models trains on the time-binned spike counts, the distribution of inter-spike intervals (ISI), or first spike latency (FSL) in individual cells (blue points) compared to shuffled data (orange points). ***B-C,*** Same as *A*, for speed and frequency range. ***D,*** Correlation between decoding accuracy in models trained on time-binned spike counts or ISIs for direction in one-octave FM sweeps. ***E-F,*** Same as *D,* for speed and frequency range. ***G-L,*** Same as *A-F,* for two-octave sweeps.

However, it is possible that there is still diversity in coding strategies used by individual neurons to encode different features—for example, some neurons may encode frequency range using the time-binned spike counts while some may encode it using the ISI distribution. To examine this, we correlated the decoding accuracies from each strategy in individual neurons. Interestingly, in the one octave dataset we found that there was no correlation between decoding strategies for sweep direction (*r^2^* = 0.002, *p* = 0.81, **Figure 5D**), speed (*r^2^* = 0.001, *p* = 0.86, **Figure 5E**), or frequency range (*r^2^* = 0.033, *p* = 0.33, **Figure 5F**). This suggests that neurons tend to favor one strategy to encode a single sweep feature rather than redundantly encoding the same information across both the time-binned spike counts and ISI distribution. In line with this, we found neurons in our dataset that encoded information using the time-binned spike counts and others that encoded information in the ISI distribution.

In contrast to results from the one-octave dataset, decoding accuracies from models trained on the time-binned spike counts were correlated with models trained on the ISI distribution in the two-octave dataset for direction (*r^2^* = 0.63, *p* = 1.4e-7, **Figure 5J**), speed (*r^2^* = 0.67, *p* = 3.4e-8, **Figure 5K**), and frequency range (*r^2^* = 0.77, *p* = 1.5e-10, **Figure 5L**). This difference in decoding strategies for one- and two-octave data may suggest differences in how neurons encode FM sweep information over time: given the shorter one-octave stimulus, information about sweep features may be present in only one spike train metric, whereas the longer two-octave stimulus may yield redundancy in coding strategies over the time course of the sound. Overall, these data suggest that IC neurons multiplex information about FM sweeps using different strategies that include the time-binned spike counts, ISI distribution, and first spike latency.

### Spike timing is important for encoding sweep speed but not sweep frequency for one- and two-octave FM sweeps

We showed above that IC neurons encode sweep direction for four-octave sweeps in a timing-dependent manner, such that disrupting the temporal information contained in the spiking pattern while leaving the overall number of spikes intact significantly decreased the direction decoding accuracy (**Figure 1K,L**). In addition, training the SVM on neuron time-binned spike counts provided the highest accuracy in decoding the frequency range and speed of the sweep in most neurons in the one- and two-octave datasets (**Figure 5**), suggesting that IC neurons may also use spike timing to encode information about these features. To directly test whether the timing of the spikes is critical for encoding these features, we trained separate SVMs to classify sound features from the time-binned spike counts for one- and two-octave FM sweeps. We then compared performance of the decoder with the time bins intact vs. with the time bins for each trial in a shuffled order, which disrupted the temporal information in each trial while preserving the total number of spikes.

We found that shuffling the order of the time bins in the one-octave condition significantly decreased the accuracy of the model for sweep direction (**Figure 6A**, paired t-test, *t* = 3.93, *p* = 0.0005), speed (**Figure 6B**, *t* = 4.61, *p* = 7.0e-5), and frequency range (**Figure 6C**, *t* = 5.62, *p* = 4.04e-6). In line with this, there were also fewer cells from which direction and speed could be significantly decoded when the time bins were shuffled (direction: 0 shuffled vs. 2 unshuffled, speed: 13 shuffled vs. 25 unshuffled), but the number of cells that encoded frequency range significantly above chance changed only minimally when the bins were shuffled (29 shuffled vs. 31 unshuffled), even though mean decoding accuracy decreased. We found a similar result in the two-octave condition, where shuffling the order of the time bins also significantly decreased the accuracy of the model for direction (**Figure 6D**, *t* = 5.19, *p* = 1.51e-5), speed (**Figure 6E**, *t* = 6.59, *p* = 3.22e-7), and frequency range (**Figure 6F**, *t* = 5.46, *p* = 7.08e-6).

**Figure 6.**
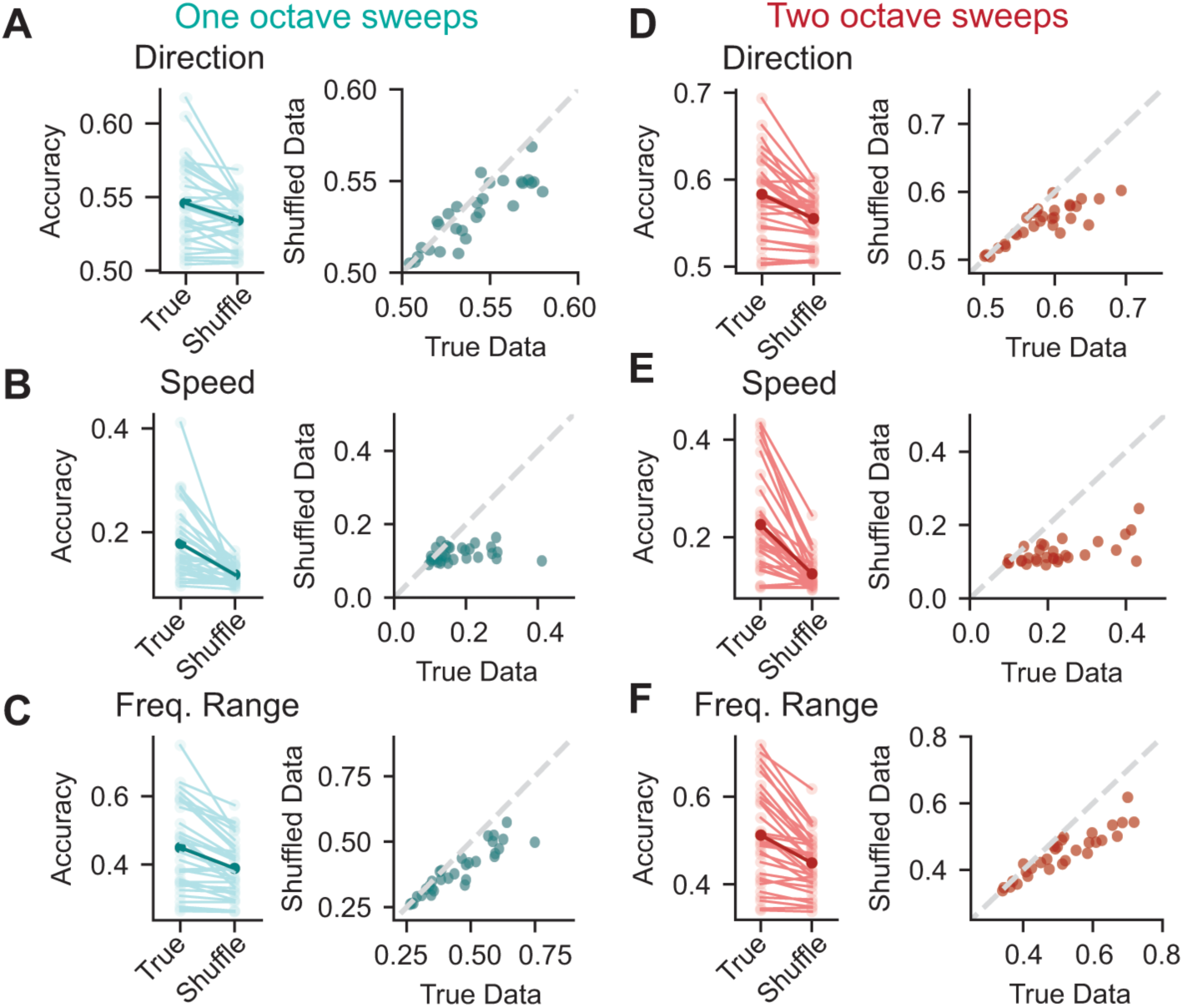
Spike timing is a critical component of feature encoding for one- and two-octave FM sweeps. *A,* Left panel: comparison of accuracy scores for decoding sweep direction for one-octave FM sweeps across each neuron where the bin order is intact (True) and shuffled (Shuffle). Class labels are maintained by trial in both cases. The mean is shown in dark blue, and light blue lines represent individual neurons. Right panel: scatterplot comparing accuracy with true bin order and shuffled bin order in individual neurons. The line of identity is shown in dashed grey (x = y). ***B-C,*** same as *A* for speed and frequency range encoding across true and shuffled bin data for one-octave FM sweeps. ***D-F***, same as *A-C* for two-octave FM sweeps.

Again, the number of cells that encoded speed and direction above chance decreased in the shuffled condition (direction: 1 shuffled vs. 9 unshuffled, speed: 11 shuffled vs. 27 unshuffled). However, despite a decrease in accuracy in the shuffled condition, the number of cells that encoded frequency range above chance remained the same in both conditions (24 shuffled vs. 24 unshuffled).

These results suggest that spike timing may be more important for conveying sweep speed compared to sweep frequency range, as frequency range decoding accuracy was less affected by disruptions in spike timing. These findings further support the idea that IC neurons can use different strategies to encode different sound features.

### IC neurons encode sound features across multiple time scales

In the analyses above, decoding algorithms were trained on all the spiking information from a neuron across an individual trial, such as the time-binned spike counts or distribution of inter-spike intervals across hundreds of milliseconds. However, postsynaptic targets are constrained in their ability to integrate inputs across time by the passive and active membrane properties of the target neurons. While this could be seen a coding limitation, this constraint would allow presynaptic neurons to encode different sound features at different times during the response, which could then be read out by one or multiple postsynaptic neurons with varying temporal integration windows.

To investigate this question, we trained an SVM to decode the direction, speed, and frequency range of an FM sweep using consecutive 6.9 ms windows (3 combined time bins) of each neuron’s response to one-octave FM sweeps of varying directions, speeds, and frequency ranges. We found that direction decoding was relatively poor throughout the duration of the sound but peaked at the beginning of the stimulus (**Figure 7A**), an effect likely due to strong onset spiking for one sweep direction in some cells. Decoding accuracy for sweep speed occurred later compared to stimulus onset but remained high throughout the duration of the sound (**Figure 7B**), whereas frequency range decoding increased rapidly after the onset of sound and then decayed (**Figure 7C**, compared in **Figure 7D**). Direction decoding was better for the two-octave sweeps compared to the one-octave sweeps and peaked at the stimulus onset (**Figure 7E**). Decoding across time was similar for the other features of the two-octave FM sweeps, with consistent decoding of speed throughout the duration of the stimulus (**Figure 7F**) and a fast increase in decoding accuracy for frequency range (**Figure 7G**) at the beginning of the stimulus.

**Figure 7.**
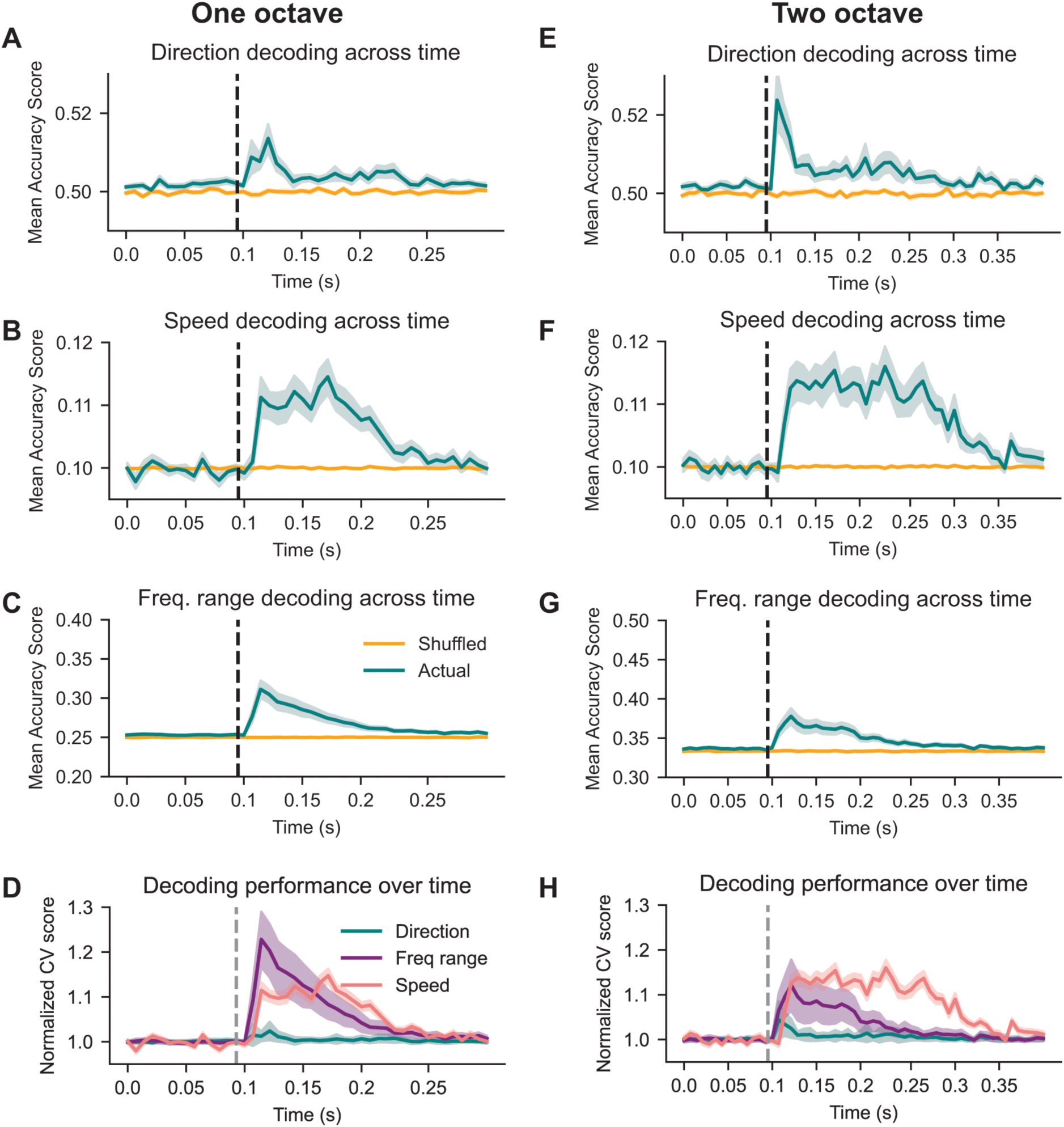
IC neurons encode FM sweep features over different time scales relative to stimulus onset. *A,* Direction decoding accuracy across time (6.9 ms time bins) for one-octave FM sweeps. The blue line indicates decoding accuracy for data with true labels, and the orange line indicates decoding accuracy for data with shuffled labels. Shading around each line represents the standard error of the mean. The dashed black line marks stimulus onset (100 ms). ***B-C,*** Same as *A* with speed and frequency range decoding across time for one-octave FM sweeps. ***D,*** Overlay of accuracy scores from *A-C* normalized to their relative pre-stimulus values. ***E-G,*** Same as *A-C* for two-octave stimuli. ***H,*** Same as *D* for two-octave stimuli.

We next asked how the accuracy for encoding each feature differed in time compared to chance level. To do this, we normalized each accuracy score to the baseline pre-stimulus accuracy (at chance level) and compared decoding accuracy over time for each feature. For both one- and two-octave sweeps, we found that accuracy for frequency range increased first and then declined, while speed decoding increased later than frequency range but was consistent throughout the duration of the stimulus (**Figure 7D,H**, pink and purple lines). Relative to chance, decoding accuracy for direction was poor across all time bins compared to the accuracy values achieved for frequency range and speed (**Figure 7D,H** blue lines).

These results suggest that individual IC neurons can represent sound features across varying time scales, which could provide another mechanism for downstream cells to receive information about complex stimuli.

### Populations of IC neurons provide robust representations of FM sweep features

While we found that decoding sweep frequency range and speed from individual neurons was above chance in many cells, accuracy values were rarely high enough to provide high-confidence information about the stimulus (**Figure 3**). However, the response heterogeneity and multiple coding strategies used by individual IC neurons could comprise a population code for various sound features. Population representations of other sound features, including sound location and amplitude-modulated stimuli, have been previously shown in IC neurons (Zohar et al., 2013; Day and Delgutte, 2016; Rogalla et al., 2024; Shi et al., 2024), but population codes for FM sweep features have not been investigated. We examined the population benefit for encoding the speed, frequency range, and direction of one- and two-octave FM sweeps by grouping cells into pseudo-populations that consisted of 2-21 neurons and compared the decoding accuracy for these groups to the accuracy obtained from single neurons.

In the one-octave dataset, we found that increasing the number of neurons in the pseudo-population (starting with decoding from individual cells) significantly increased the decoding accuracy for direction (**Figure 8A***, F*(20,630) = 58.85, *p* < 0.0001), speed (**Figure 8B** repeated measures ANOVA, *F*(20,630) = 141.8, *p* < 0.0001), and frequency range (**Figure 8C**, *F*(20,630) = 263.64, *p* < 0.0001). As cells were added to the population, direction decoding accuracy reached an asymptote that approached 65% accuracy when the population contained approximately 15 cells (**Figure 8A**). Speed decoding accuracy increased nearly linearly with each cell added to the population, and decoding accuracy was limited by the number of neurons in our dataset and thus the number of cells that could be added to the population (**Figure 8B**). Strikingly, frequency range decoding approached 100% accuracy and did not significantly increase after the pseudo-population contained 16 cells, suggesting this population size could reliably convey frequency range information for one-octave FM sweeps (**Figure 8C**).

**Figure 8.**
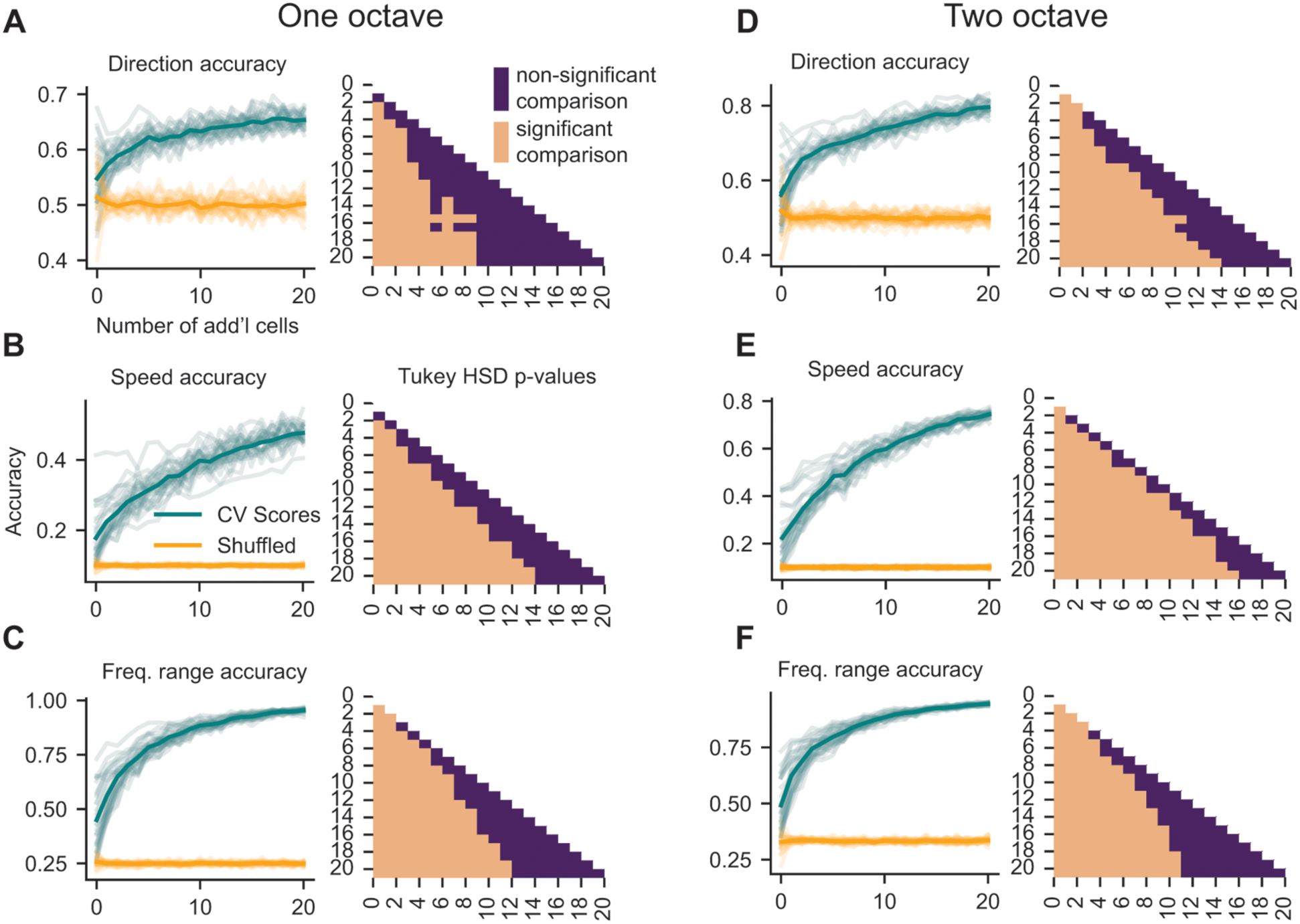
Pseudo-populations of IC neurons provide higher decoding accuracy compared to individual neurons. *A,* Left: Direction decoding accuracy of each neuron in the one-octave dataset paired with 1-20 additional neurons. Each light blue line represents decoding from one seed neuron, and the dark blue line represents the mean at each pseudo-population size. Orange lines show accuracy scores from decoding with class labels shuffled. Right: Tukey’s HSD post-hoc test corrected p-values from pairwise comparisons between the true accuracy of each pseudo-population size. Light squares represent significant comparisons while dark squares indicate comparisons that were not significantly different from each other. ***B-C,*** Same as *A* for speed and frequency range decoding. ***D-F,*** Same as *A-C* for two-octave FM sweeps.

Population decoding accuracies from the two-octave datasets largely mirrored those from the one-octave results. Increasing the number of cells in the pseudo-population significantly increased the decoding accuracy for direction (**Figure 8D**, *F*(20, 609) = 137.74, *p* < 0.0001), speed (**Figure 8E**, *F*(20, 609) = 452.70, *p* < 0.0001), and frequency range (**Figure 8F**, *F*(20, 609) = 290.97, *p* < 0.0001). Again, direction decoding accuracy was better in the two-octave dataset compared to the one-octave dataset, and decoding accuracy began to plateau around 15 cells at a decoding accuracy of 80% (**Figure 8D**). Speed decoding accuracy increased with the number of neurons in our dataset, and the trend in the data suggests that increasing the pseudo-population size would have continued to increase decoding accuracy (**Figure 8E**). In addition, frequency range decoding again approached 100% at a pseudo-population size of approximately 16 neurons (**Figure 8F**).

Overall, these results show that population codes convey more accurate information about the features of FM sweeps than individual IC neurons.

### Frequency change direction for vocalizations is not encoded in individual IC neurons

Both human speech and mouse vocalizations contain rapid frequency sweeps (Portfors, 2007), and previous studies have shown that some IC neurons fire selectively for specific vocalizations (Portfors et al., 2009; Mayko et al., 2012; Lawlor et al., 2025). However, whether the frequency sweeps present in conspecific vocalizations shape individual neuron selectivity is not well understood. To examine this, we recorded from IC neurons while playing four types of calls (calls from Schiavo et al., 2020; Valtcheva et al., 2023): an ultrasonic upsweep, an ultrasonic downsweep, an ultrasonic vocalization with complex (up and down) frequency changes, and a call with multiple low-frequency bands (in order, we refer to these as upsweep, downsweep, complex, and wriggle, **Figure 9A**).

**Figure 9.**
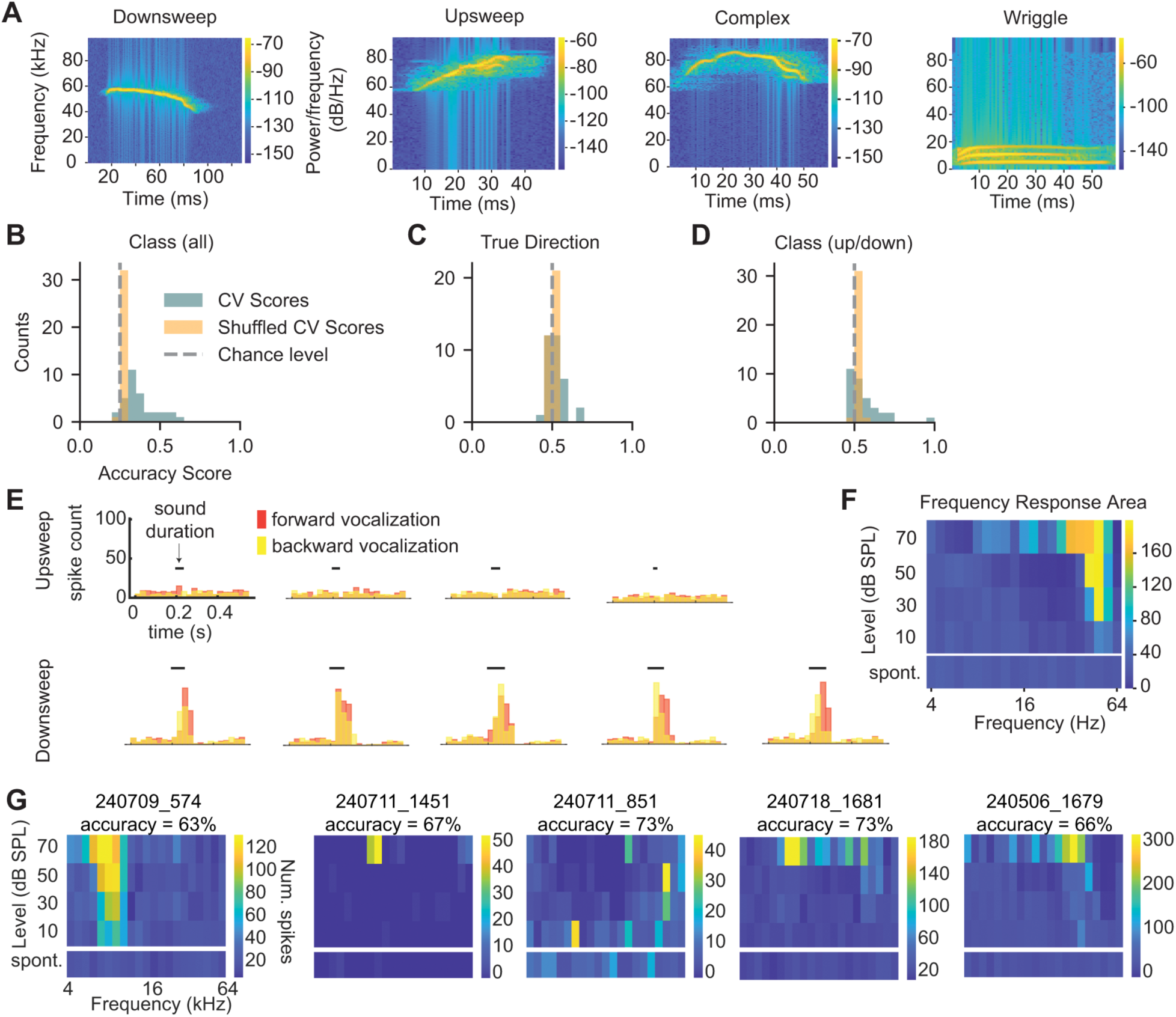
IC neuron responses to vocalizations do not depend on frequency sweep direction. *A,* Spectrograms of four example vocalizations (one from each class) used as stimuli. ***B,*** Decoding accuracy for vocalization class in individual IC neurons (blue) compared to decoding from shuffled classes (orange). Chance is labeled with a grey dashed line at 25%. ***C,*** Decoding accuracy for the true direction of the frequency sweep contained in the vocalization. ***D,*** Decoding accuracy for the vocalization class regardless of whether the stimulus was played forward or backward. ***E,*** Example peristimulus time histograms for the response from one neuron to each of the example upsweep vocalizations and downsweep vocalizations. Response to the forward vocalization is in pink and response to the backward vocalization is in yellow. The black bars indicate the sound presentation window. ***F,*** The frequency response area for the neuron in *E,* constructed from responses to 5 presentations each of 200 ms pure tones at 4 intensities and over 4 octaves with 5 frequency steps/octave. Spontaneous firing is shown in the bottom row. ***G,*** Frequency response areas from neurons where vocalization type (up/down) could be decoded significantly above chance.

Consistent with previous studies, we found that some neurons exhibited selective responses for specific vocalization classes. To quantify how well individual neurons encoded vocalization class, we trained an SVM on the spike trains of individual neurons. The SVM predicted the vocalization class above chance (25%) in 17/33 cells (**Figure 9B**), with an average accuracy of 44% in cells with significant encoding. To investigate whether the direction of the FM sweep within the vocalization was important for vocalization discriminability, we next performed decoding on two sets of responses. For the first model, we grouped responses for vocalizations where there was a true upsweep (upsweeps, and downsweeps played backwards) and trained the decoder to discriminate these from responses where there was a true downsweep (downsweeps, and upsweeps played backwards). Interestingly, we found that in only 3/33 cells in our dataset could the direction of the vocalization be decoded significantly above chance (**Figure 9C**), and average accuracy for cells with significant decoding was 62%, suggesting that most IC neurons do not specifically encode the direction of frequency sweeps contained within vocalizations.

We next asked whether the upsweep or downsweep vocalization class could be decoded using the neuron time-binned spike counts regardless of whether the vocalization was played forwards or backwards. Here, we found that 7/33 neurons could discriminate the class of the sweep, and these neurons had higher accuracy (72%) compared to the decoding for the true direction (**Figure 9D**). We examined the peri-stimulus time histograms (PSTHs) for the upsweep and downsweep vocalizations in the neuron with the strongest decoding accuracy and found exclusive responses to the downsweep vocalization regardless of what direction it was played (**Figure 9E**), indicating that information other than the direction of the frequency change is important for vocalization class selectivity in this neuron. In addition, we found that in this highly selective neuron, the tonal receptive field overlapped with the frequency content of the preferred vocalization (**Figure 9F**). However, this was not always the case: we obtained frequency response areas (FRAs) from 6/7 neurons where vocalization class (up/down) could be decoded significantly above chance and found that only two of them exhibited clear V-shaped tuning. In one of these cells (240709_574), the preferred frequency was around 10 kHz, which was much lower than the frequencies present in either vocalization (**Figure 9G**). The FRAs from the remaining four neurons exhibited some tuning at higher intensities, but again tuning did not overlap with the vocalization spectral content. These results indicate that vocalization selectivity in the mouse IC cannot be readily extrapolated from simple tonal or directional receptive fields.

## Discussion

Here, we used machine learning models to examine how multiple sound features of an FM sweep, including the frequency range, speed, and direction, are encoded in the spike trains of individual IC neurons. We found that single IC neurons can multiplex FM sweep features by encoding information along multiple dimensions of the spike train, including the time-binned spike counts, mean firing rate, ISI distribution, and first spike latency. In addition, we showed that IC neurons can encode sound features at different time points during a stimulus, providing an additional mechanism for multiplexing. We also examined decoding in pseudo-populations of IC neurons and found that, when combined, the heterogeneous responses of single neurons can yield robust population encoding of FM sweep sound features. Lastly, we showed that the directionality of frequency sweeps has little influence on vocalization selectivity in single IC neurons, indicating that complex sound responses may not be well-predicted by the receptive fields generated in individual neurons for simple sound features.

### Machine learning methods capture alternative coding strategies used by IC neurons

Many previous studies in the IC have investigated neuron selectivity for FM sweep features using selectivity indices such as the DSI, a normalized measure that compares the total number of spikes elicited by an up sweep to those elicited by a down sweep (e.g., Andoni et al., 2007; Williams and Fuzessery, 2010; Kuo and Wu, 2012; Geis and Borst, 2013). Tonal receptive fields that describe frequency selectivity in individual IC neurons are also typically constructed by comparing the total number of spikes elicited by different sound frequencies. However, total spike counts are only one way that neurons may encode sound features, and as sounds often have rich temporal structures, evaluating selectivity using only summed spike counts may omit temporal and other cues that could contain stimulus-encoding information. Indeed, emerging evidence suggests that temporal coding strategies are important not just at early auditory processing centers such as the cochlear nucleus, but also within higher-order auditory processing centers including auditory cortex (Bagur et al., 2025). Evaluating neuron encoding using only changes in spike rate also assumes that downstream neurons are acting only as simple linear summators, but a plethora of electrophysiological evidence and computational modeling suggests that this is not the case. For example, dendrites alone have numerous mechanisms that support non-linear integration (Tamás et al., 2002; Gómez González et al., 2011; see Palmer, 2014 for a review).

Here, we instead used an SVM to predict sound features from the neuron time-binned spike counts, giving the model access to firing rate across time. We found that the direction of four-octave FM sweeps could be decoded from the time-binned spike counts of many more cells than would be predicted from the DSI. In addition, we showed that disrupting spike timing while maintaining the firing rate for each trial significantly reduced the model decoding accuracy for direction in four-octave sweeps. It also significantly reduced the decoding accuracy for direction, sweep speed, and frequency range in one- and two-octave sweeps. These results indicate that there is information conveyed in spike timing about not only the direction of FM sweeps but also other sweep features, and this information is lost using methods that disregard spike timing.

### IC neurons multiplex sound features using several strategies

In this study, we highlight several temporal strategies that IC neurons use to encode features of FM sweeps, including the time-binned spike counts, the ISI distribution, and the first-spike latency (**Figure 5**). Previous studies in auditory cortex suggest that individual neurons can use some of these strategies independently of each other (Insanally et al., 2019), providing a mechanism by which neurons can multiplex features of sounds. Similarly, we found that IC neurons multiplex features of FM sweeps, including the frequency range and speed (**Figure 4**). These data are supported by other studies demonstrating joint encoding in IC neurons, for example, with encoding of sound localization cues (Chase and Young, 2008), mixed representations of sound and task variables during a behavioral task (Quass et al., 2024), and dual encoding of auditory and visual stimuli (Schmehl et al., 2025). Ultimately, a critical step toward understanding how IC neurons encode information will be to compare the encoding strategies used in the IC and the decoding strategies used by postsynaptic targets, especially in the medial geniculate body (MGB), the primary ascending target of the IC. To date, there have been few studies investigating how MGB neurons decode information from the IC, and this is an area of future research that is critical for understanding how sound information is represented along the ascending auditory pathway.

### IC neuron responses to complex stimuli are not simple summations of selectivity profiles

A common approach in past studies of the IC has been to categorize IC neuron responses by their selectivity for (or tuning to) specific sound features. However, emerging evidence from analyses of large neural datasets suggests that neuron responses are rarely truly categorical (Raposo et al., 2014; Posani et al., 2024). Here, we found that individual IC neuron responses to sweep direction varied with sweep speed and intensity (**Figure 2**). Such context-dependent selectivity has been previously shown for sound and task-dependent variables in some IC neurons (Quass et al., 2024) and for features of FM sweeps (Andoni et al., 2007; Andoni and Pollak, 2011). In this study, we also found that even in neurons that encoded FM sweep direction, this alone did not inform how those neurons responded to a vocalization with similar directional changes (**Figure 9**). However, neuron selectivity for specific sound features may not be necessary for and might even hinder robust population codes for stimuli within the IC. For example, in neocortex, a recent study suggests that categorical responses typically underlie low-dimensional rather than high-dimensional population responses (Posani et al., 2024). Similarly, studies in the IC suggest that neuron population codes for amplitude-modulated stimuli do not depend on neurons with high selectivity (Shi et al., 2024), and correlations generated between categorical neurons constrain rather than enhance the amount of information that can be contained in IC neuron populations (Zohar et al., 2013). Along the ascending auditory pathway, there is also a clear decorrelation of neuron responses to simple sound features spanning from IC to auditory cortex, suggesting that decreases in neuron selectivity parallel an increase in the specificity of the population neural code for complex auditory objects (Gosselin et al., 2025). In line with this, we found that the decoding accuracies from single neurons for sound features was on average quite low (**Figure 3**), and we did not find evidence for cells that encoded direction robustly across differing frequency ranges and speeds. However, pseudo-population decoding accuracy was high (**Figure 8**) and increased with increasing neuron numbers in the pseudo-population, pointing to robust population codes for FM sweep features in the IC that are not constrained by a lack of selectivity for features within individual IC neurons.

Overall, our data suggest that individual IC neurons can multiplex sound features using multiple strategies including both temporal and spike rate-based cues. However, most neurons do not exhibit strong spike rate-based selectivity for features of complex sounds, and neuron responses to sound features in a simple stimulus (e.g., sweep direction) do not entirely predict how the neuron will respond to natural sounds that share similar characteristics (e.g., an upward frequency sweep.) These data support an increased complexity in how neuron responses to sound features are constructed along the central auditory pathway and provide a powerful framework for understanding neural coding in the IC. An important next step for future studies is to determine how the MGB and other brain regions that receive input from the IC use the representation of complex sound features that arises from multiplexing in IC neurons.

## Materials and Methods

### Animals

All experiments were approved by the University of Michigan Institutional Animal Care and Use Committee and were in accordance with NIH guidelines for the care and use of laboratory animals. Animals were given continuous access to food and water and stayed on a 12-hour day/night cycle. Mice of both sexes were used, and all mice were on a CBA/CaJ or CBA/CaJ x C57BL/6J background to avoid the early-onset, age-related hearing loss caused by the recessive *Cdh23^Ahl+^* mutation in C57BL/6J mice (Noben-Trauth et al., 2003). To further ensure that mice used did not exhibit hearing loss, mice were aged P61-P86 at the time of experiments, as mice on CBA/CaJ and CBA/CaJ x C57BL/6J backgrounds at this age range do not exhibit age-related hearing loss on auditory brainstem response tests (Kane et al., 2012). Additionally, during electrophysiological recordings, we evaluated neural thresholds for 200 ms pure tones over a frequency range of 4 to 64 kHz. All neurons that exhibited V-shaped frequency tuning curves had thresholds at 10 dB SPL, the lowest sound level tested.

### Headbar implantation

Mice were anesthetized using 1-3% isoflurane (Piramal Critical Care, # NDC 66794-017-25), and body temperature was maintained using a homeothermic heating pad placed under the mouse. To reduce post-operative pain, mice were given subcutaneous injections of carprofen (5 mg/kg, CarproJect, Henry Schein Animal Health) and buprenorphine (0.1 mg/kg). Hair on the scalp was removed using scissors, and a circular portion of scalp approximately 11-12 mm in diameter was removed to expose the skull. The IC was identified using stereotaxic coordinates (relative to lambda and in μm: 900 caudal, 1125 lateral) and marked using a surgical marker. Bupivacaine (0.2 mL) was then applied to numb the skull, the periosteum was removed, and the skull was thoroughly scored using a #11 scalpel. To begin adhesive preparation for the headbar, a thin layer of dental acrylic (C&B Metabond from Parkell, cat #s S398, S399, S371) was applied using a fine paintbrush, which was then left to cure for ten minutes. On top, a thin layer of dental acrylic resin (Henry Schein, Inc., # 1259208 and # 1250143) was applied. Dental acrylic resin was also applied to a custom-designed headbar that was then placed onto the skull. To secure the headbar to the skull, dental acrylic resin was added in layers to cover the top of the headbar, and then silicone elastomer (Smooth-On Inc. Body Double FAST) was placed in the opening over the skull and secured with a thin strip of dental acrylic resin. Mice were monitored for signs of pain or distress for one hour post-procedure and were ambulatory before being returned to the vivarium. An additional injection of carprofen was administered the day following surgery for postoperative analgesia, and animals continued to be monitored for signs of pain or distress for 7 days.

### Craniotomy surgery

On the first day of recording, mice were anesthetized using 1-3% isoflurane. Body temperature was maintained using a homeothermic heating pad placed under the mouse. To reduce post-operative pain, mice were given a subcutaneous injection of carprofen (5 mg/kg, CarproJect). Using the IC mark that was made during the headbar implantation procedure, a craniotomy was made using intervals of drilling with a micromotor drill (K.1050, Foredom Electric Co.) with a 0.5 mm burr (Fine Science Tools). Drilling began with shallow and small circles and was interrupted every ∼20 seconds with application of cold PBS to ensure brain temperature was unaffected. When the skull above the IC was thin enough, a #11 scalpel and/or 27-gauge needle was used to remove a small section of skull above the IC. For one hour post-procedure, mice were monitored for signals of pain or distress.

### In vivo electrophysiology recordings

29 mice (14 female, 15 male) aged P61-P86 at the time of first recording were used for the study. Mice began habituation to the head fixation setup 3 or more days following headbar implantation to allow for recovery. Habituation consisted of one day of handling within the mouse’s cage and 3-5 days of head fixation inside the sound isolation booth, with the duration of head fixation increasing every day until the mouse tolerated 80 minutes of head fixation. A craniotomy was then surgically drilled above the left IC using the procedures outlined above under “Craniotomy surgery”. After one hour of recovery, mice were placed in the head fixation setup on a floating air table. The recording pipette was backfilled with an in vivo internal solution (in mM, 135 NaCl, 5.4 KCl, 1.5 CaCl_2_, 1 MgCl_2_, 5 HEPES, pH set to 7.3 with NaOH). The recording was grounded to a silver chloride wire placed into a well of PBS surrounding the craniotomy on the mouse skull. An MP-285 micromanipulator (Sutter Instrument) was used to slowly advance the pipette into the IC, and neurons were identified for juxtacellular patch-clamp recordings by an increase in electrode resistance and the presence of action potentials. Once a loose seal was made with a cell (6-47 MΩ), a regularly calibrated, free-field electrostatic speaker (Tucker-Davis Technologies, ES1) was used to present sound stimuli to the mice. The speaker was positioned contralateral to the IC recording site, at ∼10 cm in front of the right ear, at a 45° angle. Sound stimuli consisted of logarithmic FM sweeps of varying sweeps, speeds, intensities, and octaves (see Table 1), 100 ms white noise bursts (4-64 kHz, 70 dB SPL), and various mouse vocalizations presented forwards and backwards. All stimuli except vocalizations were generated in MATLAB, sampled at 192 kHz, and amplified using a TDT SA2 stereo amplifier. Vocalizations were also sampled at 192 kHz and amplified using a TDT SA2 stereo amplifier. Data were acquired at 50 kHz and low-pass filtered at 10 kHz using a Dagan BVC-700A amplifier, an NI PCIe-6343 data acquisition board, and the WaveSurfer (Janelia) package in MATLAB (Mathworks, 2022a-2024a). Individual mice were recorded from once daily for fewer than 10 sessions total. Recording locations were verified to be within the IC through post-hoc histology to visualize the electrode penetration tract, which was evident due to tissue damage and/or labeling with the fluorescent marker DiI (ThermoFisher Scientific, #V22885).

**Table 1.** Stimuli used for in vivo electrophysiological recordings.

| Stimuli | Frequency range (kHz) | Intensity (dB SPL) | Speeds (octaves/sec) | Repeats |
| --- | --- | --- | --- | --- |
| 1 octave FM sweeps | 4-8<br>8-16<br>16-32<br>32-64 | 70 | 10-200 | 10 |
| 2 octave FM sweeps | 4-16<br>8-32<br>16-64 | 70 | 10-200 | 10 |
| 4 octave FM sweeps | 4-64 at 5 steps/octave | 10, 30, 50, 70 | 10-200 | 5 |
| Pure tones (200 ms) | 4-64 at 5 steps/octave | -100, 10, 30, 50, 70 | N/A | 5 |
| Vocalizations | N/A | 70 dB | N/A | 10 |

All FM sweeps are logarithmic.

### Analysis of electrophysiological recordings

Data were re-associated with stimuli information and metadata post-collection using a custom MATLAB script. Spikes were detected using a threshold crossing approach. Detected spikes were visually examined to remove any instances where motion or other artifacts were incorrectly detected as spikes. Spike detection was further validated by performing k-means clustering on the amplitude and full-width at half-maximum amplitude of each spike; recordings that exhibited multiple clusters using this analysis were very rare and were excluded from the dataset when they were found.

### Machine learning models

Analyses were performed in Python 3.11.9. After data pre-processing (outlined above), electrophysiology data were saved into a structure and imported to Python. Support vector machine (SVM) models were implemented in Python using the ‘SVC’ function with a linear kernel from the ‘sklearn’ package. When classification was performed on more than two groups, a ‘one versus rest’ strategy was used. The C parameter was set to 1 for all models. The model features were the time-binned spike counts from each trial, and the number of bins was determined by examining model performance across 1-501 bins (steps of 2) for decoding direction of the FM sweep. The resulting bin sizes are shown in Table 2. To maintain consistency across each model, we used the bin size that resulted in the highest accuracy (2.3 ms) as features for each model. The resulting number of bins for each stimulus is shown in Table 2. The total time binned for each stimulus corresponded to the longest duration FM sweep for that stimulus set plus 20% of the stimulus time to account for time delay for sound to reach the IC and any offset spiking that might contribute to coding.

**Table 2.** Idealized bin sizes and number of bins used for analysis.

| Stimuli | Stimulus duration (ms) | Binned duration (stimulus + 20%) | Num. bins that resulted in highest decoding accuracy | Bin size (ms) that resulted in highest decoding accuracy | Num. bins used in analysis | Bin size (ms) used in analysis |
| --- | --- | --- | --- | --- | --- | --- |
| 4 octave FM sweeps | 400 | 480 | 55 | 8.7 | 209 | 2.3 |
| 2 octave FM sweeps | 200 | 240 | 75 | 3.2 | 104 | 2.3 |
| 1 octave FM sweeps | 100 | 120 | 51 | 2.3 | 51 | 2.3 |

Bin sizes and number of bins used for analysis. We first calculated the idealized bin sizes for machine learning model. To determine the best bin size to use for the decoding model, we trained the SVM to decode direction in 1, 2, and 4 octave sweeps from neuron time-binned spike counts binned using 1-501 bins in steps of 2 bins. The listed values are the bin sizes that yielded the highest direction decoding accuracy for each stimulus. Next, we calculated the number of bins needed for each stimulus to maintain a consistent time window (2.3 ms) over which each of the trials were binned. Because the 1, 2, and 4 octave stimuli were different lengths (sweep speeds were conserved across stimuli), this resulted in a different number of bins for the 4, 2, and 1 octave FM sweep stimuli.

After binning, models were trained on 80% of the data and tested on 20% of the data using 5-fold cross validation. In cases where direct comparisons were made between the sound feature decoding accuracy scores in the same neuron (for different sound features), the data were downsampled to the sound feature with the lowest number of observations and the average k-fold cross validated accuracy from each of 100 resamples was averaged and used. For example, for the 1 octave FM sweep stimulus, we played 100 sweeps/direction and 20 sweeps/speed at each frequency range, resulting in a total of 800 trials (400 per direction, 80 per speed, 100 per frequency range). Therefore, when we performed decoding analyses for frequency range, speed, and direction in the one-octave sweep condition, we used 80 trials/feature to decode each class (direction, speed, frequency range). Plots from SVM analyses were made using the ‘matplotlib’ and ‘seaborn’ packages in Python, and accuracy scores for points on the same axes are always directly comparable (had the same number of trials/class in the model).

To determine if the decoding model performed above chance in individual neurons, accuracy scores for true data were compared to accuracy scores for the model tested and trained on data with shuffled class labels. To obtain the accuracy scores for shuffled data, class labels were shuffled 100 times and the SVM was fit to the shuffled data each time. The mean and standard deviation from the distribution of shuffled accuracies was calculated for each cell. The z score for the model accuracy was calculated with reference to the shuffled distribution, and the accuracy of the model was considered above chance if this value exceeded 1.96, (the 95% confidence interval of a standard distribution with µ = 1, σ = 1).

### Comparison of decoding from spike train features

To compare decoding accuracy for FM sweep features across varying dimensions of the spike train, we calculated the distribution of the inter-spike intervals and first spike latency for each trial. To calculate the inter-spike interval, we calculated the time between each of the spikes and binned this distribution using 100 bins, meaning the model inputs were 100 features. To calculate the first spike latency, we assumed a 5 ms delay to the IC (Felix et al., 2019) and used the first spike that occurred ≥ 5 ms after the stimulus began.

### Pseudo-population decoding

To examine how groups of IC neurons encode complex sound features, we evaluated decoding accuracy in pseudo-populations of IC neurons consisting of 2-20 cells. First, we examined coding in an individual neuron, as described above. Next, we randomly sampled 10 neurons from our population and paired each of these neurons individually with the first neuron. To evaluate decoding for neuron pairs, we horizontally concatenated the time-binned spike counts (rows) for each neuron pair for trials with matched stimuli and then evaluated the performance of the model on the concatenated data. To find the overall accuracy for the individual cell plus a paired cell, we averaged the model accuracy for each pairing. We repeated this process with every cell in our dataset, pairing it with 10 randomly selected neurons and evaluating average model accuracy across the 10 pairs to get an overall paired accuracy for the population.

We next continued this process by increasing the number of paired cells for each individual neuron the dataset. For each individual cell, there were 10 randomly selected groups of neurons that were paired (e.g., 10 groups of 2, 3, 4, 5… etc.) and decoding was performed on the concatenated time-binned spike counts from these groups. For each cell, the average accuracy from each set of pairings was calculated, and then the average accuracy across the dataset was plotted.

### Statistical tests

All statistical tests were performed using Python 3.11.9. To evaluate whether there was significant decoding accuracy in individual neurons, we z-scored accuracies relative to shuffled class accuracy distributions, as described above. To evaluate whether there were differences in decoding accuracy between true and shuffled bin conditions, data were evaluated using a paired t-test (‘ttest_rel’ function from ‘scipy.stats’). To examine correlations between decoding for sound features, we performed a linear regression.

To examine direction decoding accuracy across other varying sweep parameters such as sweep intensity and speed, a repeated-measures ANOVA (‘ANOVArm’ from ‘statsmodels.stats.anova’) was performed followed by Tukey’s HSD post-hoc tests (‘pairwise_tukeyhsd’ from ‘statsmodesl.stats.multicomp’) where appropriate. We also used a repeated-measures ANOVA and Tukey’s HSD post-hoc test to examine if increasing the number of cells in each pseudo-population increased the average accuracy. To test for an increase in the number of cells that significantly encoded direction over increasing speed or intensity, we used a Chi-square test for trend (‘chi2_contingency’ from ‘scipy.stats’), which can be used to examine changes in proportions over ordered groups.

All p-values were compared to a Bonferroni-corrected α value for evaluation of significance where appropriate, and in some cases Bonferroni-adjusted p-values are instead displayed and noted as “adj. p-value”.

### Direction selectivity index calculation

The direction selectivity index (DSI) was calculated as follows, where spikes_up_ indicates the number of spikes that occurred during upward frequency sweeps and spikes_down_ indicates the number of spikes that occurred during downward frequency sweeps:

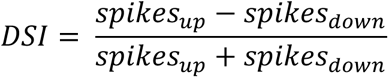

The time window across which spikes were summed for the 4-octave frequency sweeps was 480 ms after stimulus onset (since the longest stimulus was 400 ms, this is stimulus time + 20% of stimulus time to account for any offset spiking that may be directionally selective). This is the same time window over which spike times were binned for the machine learning analysis.

## Data Availability Statement

All data will be made available upon reasonable request to the corresponding author.

## Acknowledgements

The authors thank Pierre Apostolides, Laurel Carney, Yoani Herrera, and Susan Shore for helpful discussions, advice and support. The authors would also like to thank the Robert Froemke lab at New York University for providing the mouse vocalization recordings used in this study. This work was supported by a Rackham Graduate Student Research Grant (ACD), the National Science Foundation Graduate Research Fellowship Program under Grant No. DGE 1256260 (ACD), the National Institutes of Health Grant R01 DC018284 (MTR), the National Institutes of Health Grant F31 DC021344 (ACD), and the National Institutes of Health Grant K99/R00 DC019415 (MAS).

## Author contributions

ACD, SZW, MAS, and MTR conceived of the study and designed experiments. ACD, SZW, and MAS conducted experiments. ACD, SZW, MM, RW, and MTR analyzed data. ACD and SZW prepared figures. ACD and SZW prepared the initial draft of the manuscript. ACD, SZW, MM, MAS, RW and MTR revised the manuscript, and all authors approved the submitted version.

